# Interleukin-25-mediated resistance against intestinal trematodes does not depend on the generation of Th2 responses

**DOI:** 10.1101/2020.03.02.972836

**Authors:** María Álvarez-Izquierdo, Miguel Pérez-Crespo, J. Guillermo Esteban, Carla Muñoz-Antoli, Rafael Toledo

**Affiliations:** Área de Parasitología, Departamento de Farmacia y Tecnología Farmacéutica y Parasitología, Facultad de Farmacia, Universitat de València, Avda. Vicent Andrés Estellés s/n, 46100 Burjassot, Valencia, Spain

## Abstract

Interleukin-25 (IL-25) is recognized as the most relevant initiator of protective Th2 responses in intestinal helminth infections. It is well known that IL-25 induces resistance against several species of intestinal helminths, including the trematode *Echinostoma caproni*. *E. caproni* has been extensively used as an experimental model to study the factors determining the resistance to intestinal infections. Herein, we assessed the role of IL-25 in the generation of resistance in mice to *E. caproni* infections. ICR mice are permissive hosts for *E. caproni* in which chronic infections are developed in relation to the lack of IL-25 production in response to primary infection and the consequent development of a Th1 response. However, pharmacological clearance of the primary infection induces non-specific expression of IL-25 that protects mice to secondary challenge infections in association with Th2 responses. Using this experimental model, we have determined that the role of IL-25 in the polarization of the immune response differs between the primary and secondary memory response. IL-25 is required for the development of a Th2 phenotype in primary *E. caproni* infections but also promotes the differentiation to Th2 memory cell subsets that enhances type 2 responses in memory responses, even in the absence of IL-25. Despite these events, development of Th2 responses does not induce resistance to infection. Our results suggest that Th2 phenotype does not elicit resistance and IL-25 is responsible for the resistance regardless of the type 2 cytokine activity and STAT6 activation. Alternative activation of macrophages induced by IL-25 could be implicated in the resistance to infection. In view of the critical role of IL-25, we have also investigated the factors determining the production of IL-25 and appears to be related to the alterations in resident microbiota induced by the infection.

**Author’s summary:** Interleukin-25 (IL-25) plays a major role in resistance against intestinal helminth infections as initiator of protective Th2 responses. However, recent studies have challenged the contribution of this cytokine in both the polarization of the response towards a Th2 phenotype and the parasite rejection. We have used the experimental model *Echinostoma caproni*-ICR mice to investigate the participation of this cytokine in resistance to intestinal helminths. ICR mice are characterized by their inability to respond with IL-25 production in primary infections with *E. caproni*, causing susceptibility associated with a Th1 response. However, mice are refractory to infection in presence of IL-25 in relation to a type 2 phenotype. Herein, we show that dynamics of resident microbiota appears to be crucial in IL-25 production. Moreover, IL-25 seems to play a pivotal role in the polarization to Th2 in primary responses, but also appears to participate in the generation of memory mechanisms making unnecessary the participation of IL-25 in memory responses for the development of Th2 milieu. However, resistance to *E. caproni* infection does not depend on the generation of a Th2 phenotype, but exclusively depends on the presence of IL-25, operating autonomously from the type 2 response in the generation of resistance.

## Introduction

Intestinal helminth infections are common in man and animals, especially in developing regions of Africa, Asia and the Americas [1–3]. These parasitic infections generate substantial mortality and morbidity and produce relevant physical and mental disorders and often persist in the face of serious economic problems [2]. Moreover, infections by intestinal helminths also compromise the health and productivity of livestock worldwide [3]. Currently, the impact of intestinal helminth infections only can be reduced by anthelminthic treatment, but the progressive emergence of resistances to these drugs limits their utility. Furthermore, the fact that infections do not generate protective immunity causes continuous reinfections in environments of poverty and poor sanitary conditions. Despite these facts, no available effective vaccines to protect humans or animals exist. Among other factors, the lack of knowledge on how protective immunity is selected after infection is a major obstacle to successful immunization [4].

Resistance to intestinal helminths is based on the generation of Th2 responses in a complex process that involves the interaction between innate and adaptative mechanisms [5–7]. Protective Th2 immunity against intestinal helminths is initiated and amplified by the epithelial-derived alarmin cytokines including IL-25, IL-33 and thymic stromal lymphopoietin (TSLP), though the immune mechanisms behind the development of these responses are poorly understood [6, 8]. In recent years, IL-25, a member of the IL-17 family of cytokines also called IL-17E, has been considered a key cytokine. IL-25 promotes Th2 immunity and exerts anti-inflammatory functions via the downregulation of Th17 and Th1 responses [9–12]. IL-25 expression is generally associated with resistance to gastrointestinal helminth infections through the activation of Th2 responses that mediate effector mechanisms for parasite expulsion (which include goblet cell hyperplasia, smooth muscle hypercontractility, expression of RELM-β, and intestinal mastocytosis, amongst others) [6]. Recent work has uncovered the origin and the mechanisms of action of IL-25 [13–15]. Intestinal tuft cells are the main source of IL-25. Upon helminth establishment, tuft cells release IL-25. In response to alarmins, group 2 innate lymphoid cells (ILC2) produce large amounts of IL-13 that activates dendritic cells in the lamina propria and license their migration to mesenteric lymph nodes to polarize naïve CD4+ T cells into Th2. ILC2 and basophils can also perform antigen presentation to CD4+ T cells and induce Th2 polarization, which is aided by IL-4 in the case of basophils. Th2-polarized cells release an array of cytokines that drive effector mechanisms and expand themselves through positive feedback loops, amplifying the response [6]. Despite these facts, there are several doubts in relation to the role of IL-25 in the generation of protective Th2 responses to intestinal helminth infections [8,16–17]. For example, it is not well defined if the participation of IL-25 is limited to its ability to promote Th2 responses or if it is directly involved in the activation of effector mechanisms responsible for resistance. Likewise, its role in the differentiation of Th cells to memory subset cells and their implications in the generation of immunity against intestinal helminths is unknown. Several recent studies have questioned the role of IL-25 on the generation of adaptive type 2 responses or the differentiation of Th2 cells or their development to effector or memory Th2-cell subsets [8, 16].

Apart from their interest as human parasites mainly in East and Southeast Asia [18–19] echinostomes, and particularly *Echinostoma caproni* (Trematoda: Echinostomatidae), have been extensively used for the study the factors on which rely the establishment of chronic infections or, in contrast, the development of resistance to intestinal helminths. *E. caproni* is and intestinal trematode with no tissue phase in the vertebrate definitive host [20]. After infection, the metacercariae excyst in the duodenum and the juvenile worms migrate to the ileum, where they attach to the mucosa. *E. caproni* has a wide range of definitive hosts, although its compatibility differs considerably between rodent species in terms of worm survival and development [21]. In mice and other hosts of high compatibility, the infection becomes chronic, while in hosts of low compatibility, (e.g. rats) the worms are expelled from the 2-4 weeks post-infection [22–23]. The establishment of chronic infections in ICR mice is dependent upon a local Th1 response with elevated production of IFN-γ [24]. In contrast, the resistance to *E. caproni* infection in hosts of low compatibility is associated with the development of a local Th2 phenotype [24–25]. Because of these characteristics, the *E. caproni*-rodent model is useful to elucidate several aspects of the host-parasite relationships in intestinal infections, such as the induction of distinct effector mechanisms and their effectiveness in parasite clearance. Recent studies of our group showed that partial resistance against *E. caproni* secondary infections is developed after chemotherapeutic cure of a primary infection and innately produced IL-25 is crucial to determine the resistance. Susceptibility to primary infections was associated with low levels of intestinal IL-25 expression, whilst deworming via administration of praziquantel (pzq) was accompanied by a steady increase in IL-25 expression and, in turn, by the onset of a Th2-type response that prevented the establishment of secondary infections [26–27].

In the present work, we investigate the role of IL-25 in resistance to *E. caproni* infections in mice. Our results show that IL-25, but not type 2 response, is required for resistance. However, IL-25 may have a rolo in the differentiation of Th2 memory subset, facilitating Th2 responses to challenge infections. Susceptibility of mice to *E. caproni* infections relies on the fact that parasite compounds do not elicit IL-25 responses in mice and the upregulation of this cytokine depends on external factors such as changes in resident microbiota.

## Material and methods

### Parasites, hosts and experimental primary and secondary infections

The strain of *E. caproni* has been described previously [28]. Encysted metacercariae of *E. caproni* were removed from the kidney and pericardial cavity of experimentally infected *Biomphalaria glabrata* snails and used to infect male ICR mice weighing 30-35 g by gastric gavage min both primary and challenge infections (50 metacercariae each. The positivity of the infection in each case was determined at necropsy or detection of eggs in stools as described previously [22]. Animals were maintained under conventional conditions with food and water ad libitum.

### Ethical statement

This study has been approved by the Ethical Committee of Animal Welfare and Experimentation of the University of Valencia (Ref#A18348501775). Protocols adhered to Spanish (Real Decreto 53/2013) and European (2010/63/UE) regulations.

### Pharmacological treatment of primary infections

Curation of the primary infections was achieved by pharmacological treatment with praziquantel. Mice were treated with a double dose of 100 mg/Kg of pzq at 4 weeks post-primary infection (wppi), orally administered on alternate days as described previously [29]. All mice belonging treated with pzq responded to the treatment and reverted to negative as determined by coprological examination. The influence of the pharmacological treatment over the studied parameters was discarded since five mice were left uninfected, treated with pzq as described above and analyzed as the other animals.

### Treatment of mice with commercial, specific blocking antibodies or isotype-matched IgG control antibodies

Briefly, several mice were sensitized by intraperitoneal injection with commercial specific blocking antibodies, recombinant proteins or isotype matched control antibodies.

To neutralize the effect of IL-25 produced nonspecifically after the curation of the primary *E. caproni* infection and to investigate the effect of this cytokine in a secondary challenge infection and the role of STAT6 activation in resistance to infection, two group of 5 mice each were treated with monoclonal anti-mouse IL-25 (mα-IL-25) (R&D Systems) or monoclonal anti-mouse IL-4Rα (mα-IL-4Rα) (Biolegend). In each group, mice were primarily infected with 50 metacercariae of *E. caproni* and treated with pzq at 4 wppi. On each of the two days previous to a secondary infection at 6 wppi, mice were intraperitoneally injected with either mα-IL-25 (concentration: 0.25 μg/μl) in one of the groups or mα-IL-4Rα (concentration: 0.1 μg/μl) to the mice belonging to the other group in 150 μl of saline buffer. Additionally and to be used as control of the mα-IL-25-treated mice, 5 mice were injected with rat IgG1 and other 5 mice with IgG2b and used as control of the mα-IL-4Rα-treated group of animals. All mice were sacrificed at 2 weeks post-secondary infection (wpsi).

Moreover, several recombinant cytokines were used to analyze their effect on the course of the infection. For this purpose, groups of 5 mice were intraperitoneally injected with either IL-4 (rIL-4; Prepothech), IL-13 (rIL-13; Prepotech) or IL-25 (rIL-25; R&D Systems) (concentration: 0.2 μg/μl each) in 150 μl of PBS during each of the four days from the primary infection with 50 metacercariae of *E. caproni*. All mice were sacrificed at 2 wpsi. Additonally, a group of five mice were treated with recombinant IL-13Rα2 (rIL-13Rα2; R&D Systems) (concentration: 0.2 μg/μl) following identical protocol to study the role of IL-13Rα2 in *E. caproni* infection. As control animals, five mice were intraperitoneally injected with 150 μl of PBS following the same protocol. Finally, a total of 15 also were treated with rIL-25 and primarily infected as described above and at 2 post-tretament with pzq (wppt). Every two wppi, 3 of these mice were sacrificed to analyze the kinetics of endogenous IL-25 expression. Once IL-25 levels returned to baseline, the remainder 3 mice were secondarily infected with 50 metacercariae of *E. caproni*. Five other mice were infected, treated, aged and reinfected following the protocol described for the previous group and sacrificed simultaneously as a control.

### Total RNA extraction

Total RNA was extracted from full-thickness sections of ileum of necropsied mice. Total RNA was isolated using Real Total ARN Spin Plus kit (Durviz) according to the manufacturer’s instructions. The cDNA was synthesized using High Capacity cDNA Reverse Transcription kit (Applied Biosystems).

### Real-Time PCR and relative quantification analysis

For quantitative PCR, 40 ng total RNA wasreverse transcribed to cDNA and added to 10µL of TaqMan^®^ Universal PCR Master Mix, No AmpErase^®^ UNG (2x), 1µL of the specified TaqMan^®^ Gene Expression Assay, and water to a final reaction volume of 20µL. Reactions were performed on the Abi Prism 7000 (Applied Biosystems^®^), with the following thermal cycler conditions: initial setup of 10 min at 95 °C, and 40 cycles of 15 s denaturation at 95 °C and 1 min of annealing/extention at 60 °C each. Samples were amplified in a 96-well plate. In each plate, endogenous control, samples and negative controls were analyzed in triplicate. All TaqMan^®^ Gene Expression primers and probes for inducible nitric oxide synthase (iNOS), cytokines and mucins were designed by Applied Biosystems^®^ and offered as Inventoried Assays. The assay ID details are shown in Supplementary Table 1. Each assay contains two unlabeled primers and one 6-FAM™ dye-labeled, TaqMan® MGB probe. Primer concentration was optimized by a matrix of reactions testing a range of concentrations for each primer against different concentrations of the partner primer and also negative controls were included.

Cycle threshold (Ct) value was calculated for each sample, housekeeping and uninfected control. To normalize for differences in efficiency of sample extraction or cDNA synthesis we used β-actin as housekeeping gene. To estimate the influence of infection in the expression levels we used a comparative quantification method (2^-ΔΔCT^ – method)^51^. This method is based on the fact that the difference in threshold cycles (ΔCt) between the gene of interest and the housekeeping gene is proportional to the relative expression level of the gene of interest. The fold change in the target gene was normalized to β-actin and standardized to the expression at time 0 (uninfected animals) to generate a relative quantification of the expression levels.

### Analysis of goblet cell responses

Goblet cell responses to *E. caproni* infections in the ileum of mice were evaluated in primary and secondary infections in rIL-25-treated mice. At each time point, 5 mice in each group were necropsied and ileal sections of about 0.7 cm in length were obtained and fixed in 4% paraformaldehyde (PFA). After embedding in paraffin wax, serial 4 μm-sections were cut from each tissue block and stained with alcian blue.

### Indirect immunofluorescence

Translocation and phosphorylation of STAT6 were study by fluorescent immunohistochemistry performed on paraffin-embedded tissue sections basically as described previously [30]. Rabbit antibodies anti-STAT6 (ThermoFisher Scientific) and anti-p-STAT6 (ThermoFisher Scientific). Anti-STAT6 and anti-p-STAT6 were diluted 1/200 and 1/20, respectively, in PBS containing 0.3% Triton™ X-100 and 10% FCS and incubated for 2h in a humid chamber at room temperature, under continuous agitation. After washing 3 times in PBS, intestinal sections were incubated for 2 h with secondary antibody, goat anti-rabbit IgG conjugated with Alexa Fluor® 647 (Jackson ImmunoResearch Laboratories, Inc.), diluted 1/600 in PBS-Triton™ X-100 (0.3%) for Anti-STAT6 and 1/100 for anti-p-STAT6. Slides were washed in PBS and cell nuclei were counterstained with DAPI before mounting with Fluoromount™ (Sigma-Aldrich). Cell staining was analyzed by fluorescence microscopy. Results were studied over 5 selected fields.

### Enzyme immunohistochemistry with Diaminobenzimidine (DAB)

To analyze the tuft cells and GATA3+ cells such as ILC2 and Th2 populations in primary and secondary infections with *E. caproni*, enzymatic immunohistochemistry of intestinal sections was performed. Initially, intestinal samples were dewaxed by incubating them for 20 minutes in an oven at 60 ° C, passed through a hydration chain (Xylene 4x 5 min - 100% Ethanol 2x 3 min - 90% Ethanol 2x 3 min - 70% Ethanol 2x 3min) and were incubated in 10 mM Sodium Citrate Buffer for 10 min to improve antigen detection. Once the samples were cooled, they were kept in running water for 10 min and washed twice for 5 minutes in Tris Buffer Saline (TBS) + 0.1% Triton X-100 pH 7.6 while stirring. Sections were blocked with 2.5% Normal Goat Serum (Vector) for 1 hour at room temperature. After blocking, they were incubated over night at 4 ° C with the primary antibody at 1: 1000 dilution in TBS and 1% BSA. As primary antibodies were used: a-DCLK-1 (Abcam) for labeling tuft cells and a-GATA3 (Abcam) for labeling GATA3+ cells.

In order to eliminate the own autofluorescence from the tissue, the samples were incubated in Dual Endogenous Enzime Block (Dako) for 10 min at room temperature. They are then incubated with the secondary Polyclonal Goat anti-Rabbit Immunoglobulins HRP (Dako) antibody diluted 1: 1000 in TBS + 0.1% Triton X-100 and 1% BSA for 1 hour at room temperature. After all incubation steps, 2 washes of 5min were performed with TBS + 0.1% Triton X-100 with gentle agitation.

DAB was selected as a chromogen to reveal the reaction (Liquid DAB + Substrate Chromogen System, Dako). The development time must be controlled by observing the brown precipitates produced by reacting the DAB with the Ab2-HRP. The samples are washed with running tap water to stop the reaction.

Finally, the sections were contrasted with Mayer’s Hematoxilin (Dako), passed through a dehydration chain (Scott’s tap water 30 sec - 90% Ethanol 2×30sec - 100% Ethanol 2x 3min - Xylene 2x 3 min) and mounted in DPX liquid medium for later analysis in an optical microscope at a magnitude of 200x magnification. Cell populations were studied over 10 selected representative fields.

### Induction of intestinal dysbacteriosis

To analyze the effect of resident microbiota on the production of IL-25 in response to *E. caproni* infection. A total of 10 were primarily infected and treated with pzq at 2 wppi. To avoid recovering of the microbiota after curation and to evaluate its role in a secondary infection, dysbacteriosis was induced in in five of these mice using a cocktail of antibiotics. Mice received drinking water containing ampicillin (Sigma) (1.0 g/l), metronidazol (Guinama) (1.0 g/l), neomycin (Sigma) (1.0 g/l) and vancomycin (Sigma) (0.5 g/l) for two weeks before secondary infection with 50 metacercariae of *E. caproni*. The remainder 5 mice were secondarily infected at 2 wppt without antibiotic treatment. All mice were necropsied at 2 wpsi and expression of IL-25 was compared with that of secondary infections in mice not treated with antibiotics. Additionally, 5 mice were used to evaluate the potential effect of antibiotic treatment in the expression of IL-25. These mice primarily infected and treated with pzq and antibiotics following the same procedure but not secondarily infected. No changes in IL-25 were observed in these control animals.

### Determination of the total bacterial load in fecal samples

In order to determine the total bacterial load in the fecal samples, qPCR of 16S rRNA gene was performed. For this purpose, the KAPA SYBR. FAST qPCR Kit was used. For each sample, 20 μl PCR duplicates were prepared with each containing 2μl of the DNA used as template, 10μl of mix provided by the manufacturer, and 0.4μl of forward and reverse primers at the final concentration of 0.2mM (27F-qPCRAGAGTTTGATCMTGGCTCAG; 338R-qPCRTGCTGCCTCCCGTAGGAGT). In order to complete the volume of the reaction, 7.2 μl of water was added.

A PCR product of the 16S rRNA gene from *Enterococcus faecium* C68 strain was used for obtaining a standard curve. This *E. faecium* 16S rRNA PCR was performed as follows. Briefly, 25μl reaction was prepared containing 1μl of 1 bacterial colony resuspended in PBS, 2.5μl 10x Standard Taq Reaction Buffer (New England BioLabs), 0.25 mM of deoxynucleoside triphosphates (dNTPs), 2.5 U of Taq DNA Polymerase (New England BioLabs) and 0.2 mM of primers. The volume was completed with water.

ENDMEMO program was used in order to determine the number of 16S rDNA molecules in the PCR product of *E. faecium* CD68 based on sequence of 16S rRNA gene and concentration of the PCR product. A standard curve was obtained by making 5-fold dilutions of the PCR product. Cycling conditions of the qPCR were 94.C for 5 minutes, and 45 cycles of 94.C for 30 seconds, 56.C for 30 seconds and 68.C for 30 seconds, and a final elongation cycle at 68.C for 5 minutes. By extrapolation of results with the ones obtained with standard curve, the number of 16S rRNA genes was determined for each sample. The final number of 16S rRNA genes per gram of fecal sample was calculated by using the following formula:

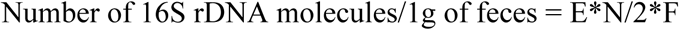

where E represents the volume of the buffer used for elution of DNA after extraction, N represents number of 16S rDNA molecules obtained by qPCR, 2 stands for the volume of DNA used for the qPCR reaction, and F represents the weight (in grams) of the fecal pellet from which DNA was extracted.

### Statistical analysis

χ^2^ test was used to compare the rates of infection between both groups of mice at each week post-infection. To compare the worm recovery between primary and challenge infections, a Student’s t-test was used at each week post-infection. One-way ANOVA with Bonferroni test as post-hoc analysis were used to compare expression levels of cytokines, enzymes, or other genes analyzed by PCR. P<0.05 was considered as significant. Prior to analyses, data were log transformed to achieve normality and verified by the Anderson–Darling Test.

## Results

### Treatment of mice with rIL-4 or rIL-13 does not induce resistance to primary infection

Susceptibility of mice to primary *E. caproni* infection has been attributed to the inability of mice to respond with overproduction of type 2 cytokines, especially IL-4 and IL-13. To analyze the role of these cytokines in the course of a primary *E. caproni* infection, we infected rIL-4- and rIL-13-treated mice and compared with an untreated group of mice (n=5). The results obtained show that neither treatment induced resistance to primary infection. The worm recovery in rIL-4-[51-65% (61.2 ± 11.2)], rIL-13-treated [62-85% (76.1 ± 16.5)] or non-treated mice [54-68% (62.1 ± 10.6)] was very similar (Fig. 1A).

**Figure 1.**
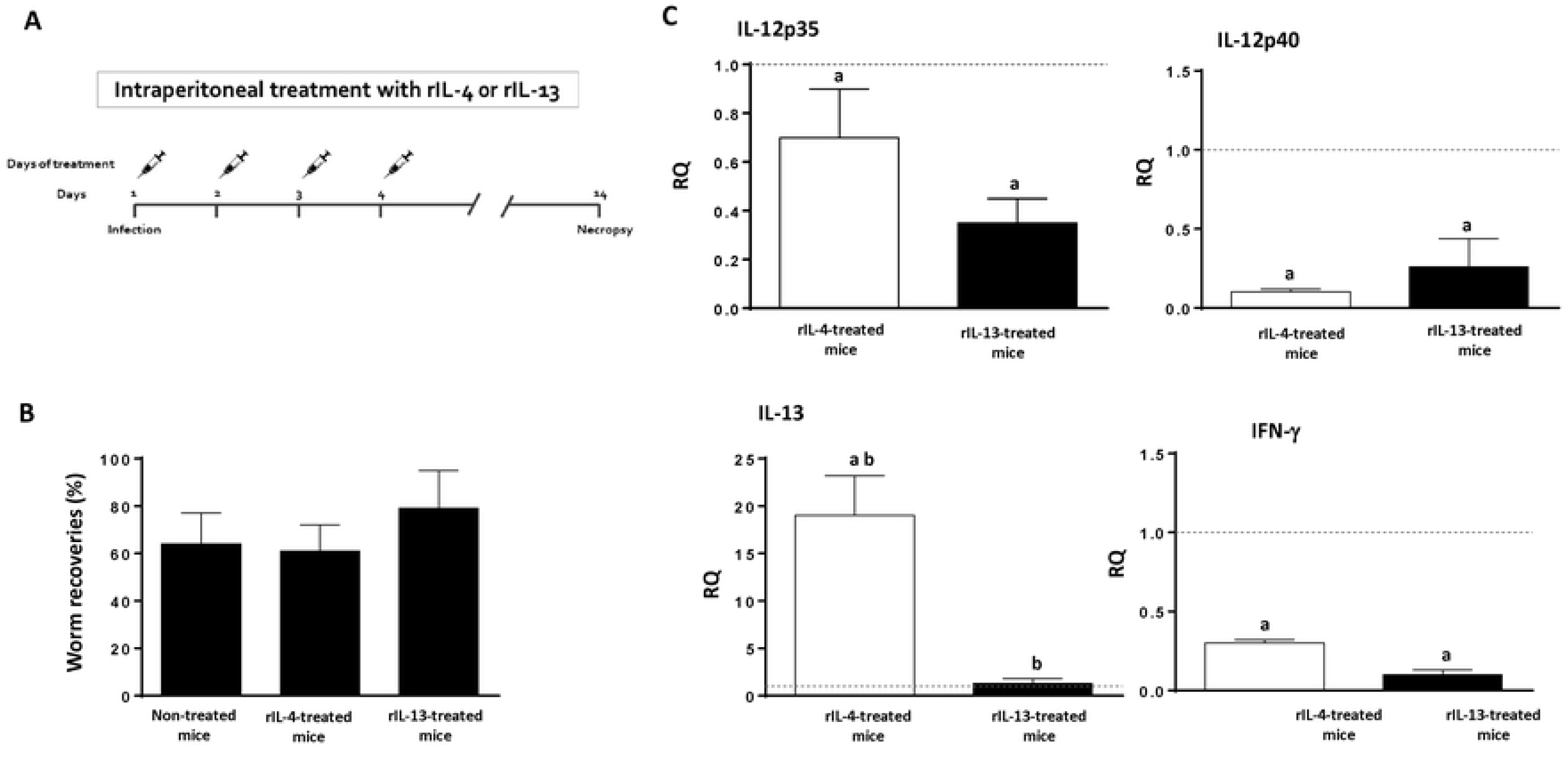
Treatment of mice with rIL-4 or rIL-13 induces a Th2 response to primary *Echinostoma caproni* infection but not resistance to infection. (A) Schematic representation of the experimental protocol; (B) Counts of tuft cell populations; (B) worm recovery in naïve, rIL-4-treated and rIL-13-treated mice mice; (C) expression of cytokine mRNA in the intestinal tissue of rIL-4-treated or rIL-13-treated mice at two weeks post-primary infection with *E. caproni*. The relative quantities (RQ) of cytokine genes are shown after normalization with β-actin and standardization of the relative amount against day 0 sample. Vertical bars represent the standard deviation. a: significant differences with respect to naïve mice controls; b: significant differences between groups (p<0.05).

Treatment of mice with rIL-4 or rIL-13 only induced slight changes in cytokine expression and only decreases in IL-4 and IL-12p35 were detected in rIL-4-treated mice. (Supplementary Fig. 1). Infection of treated mice elicited a marked downregulation of type 1 cytokines such as IL-12p35, IL-12-p40 or IFN-γ. In rIL-4-treated mice, a significant upregulation of IL-13 also was observed (Fig. 1B). No significant differences between groups were detected in the remainder cytokines (data not shown). Similarly, no changes were observed in the expression of markers of macrophage activation (data not shown).

Treatment with rIL-13 induced goblet cell hyperplasia that were more pronounced after infection of the treated mice. In contrast, only a weak hyperplasia after the experimental infection was observed in the animals treated with rIL-4 (Fig. 2A-E). RELM-β expression became downregulated in animals treated with rIL-4 but, in contrast, weak upregulation was observed in rIL-13-treated mice at 2 wppi (Fig. 2F).

**Figure 2.**
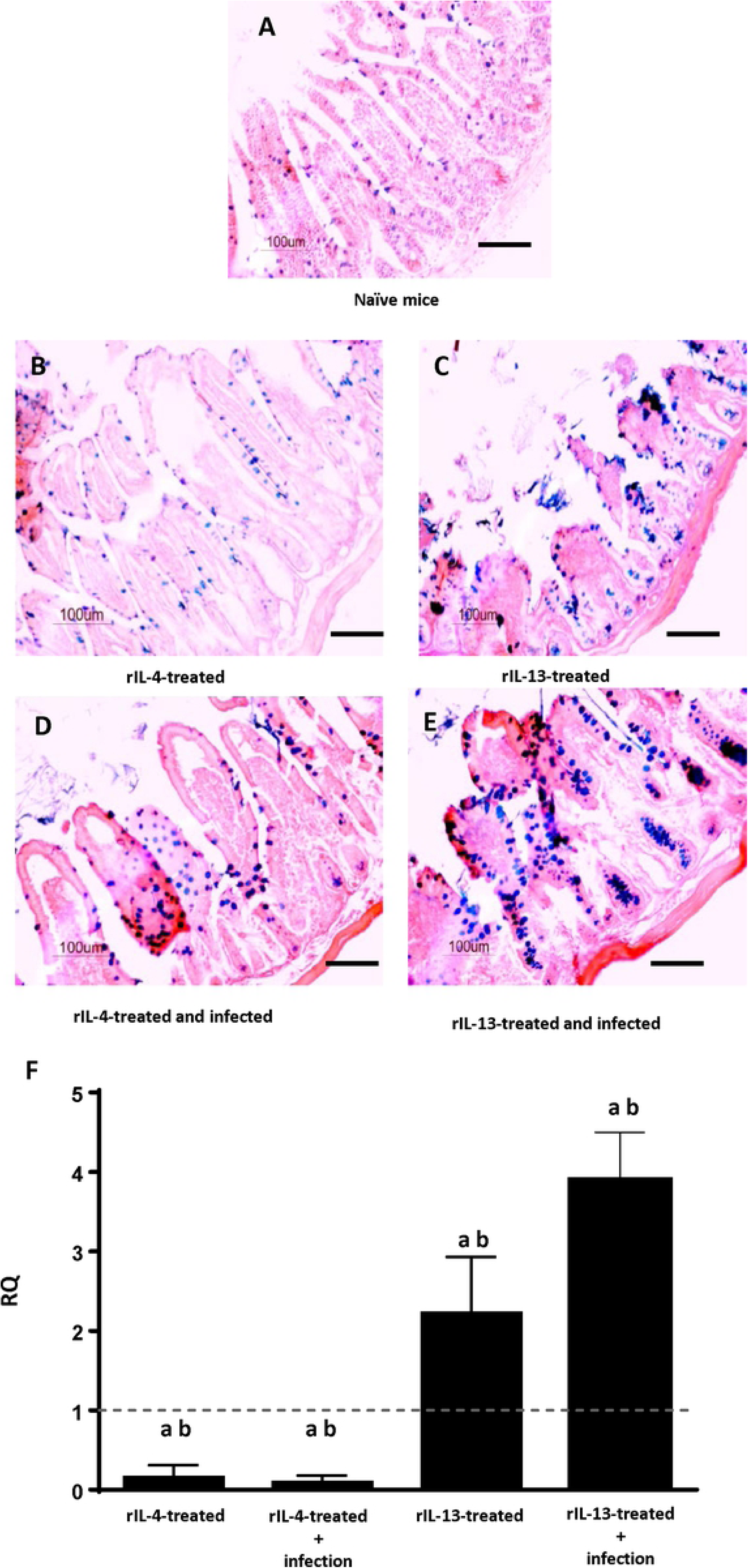
Treatment with rIL-4 or rIL-13 induces goblet cell expansion and overexpression of RELM-b after a primary infection with *Echinostoma caproni*. Alcian blue staining of intestinal tissue (A-E) and expression of RELM-β mRNA in the intestinal tissue (F) of naïve mice, non-infected rIL-4- or rIL-13-treated mice and infected rIL-4- or rIL-13-treated mice at 2 weeks post-primary infection. Vertical bars represent the standard deviation. a: significant differences with respect to negative controls; b: significant differences between groups at each week of the study (p<0.05).

### Secondary *E. caproni* infection induces expansion of tuft cells and GATA3+ cells concomitantly with a Th2 response

To evaluate the mechanisms by which IL-25 participates in resistance to E. caproni infection, we analyzed the kinetics of tuft cells and GATA3+ cells such as 2 ILC2 or Th2 in primary and secondary infections (Fig. 3 and Supplementary Fig. 2). Primary infection did not elicit hyperplasia of either tuft cells or GATA3+ cells, and no increase was observed after treatment with pzq. In contrast, a marked increase in counts of tuft cells and GATA3+ cells was observed as a consequence of the secondary infection (Fig. 3).

**Figure 3.**
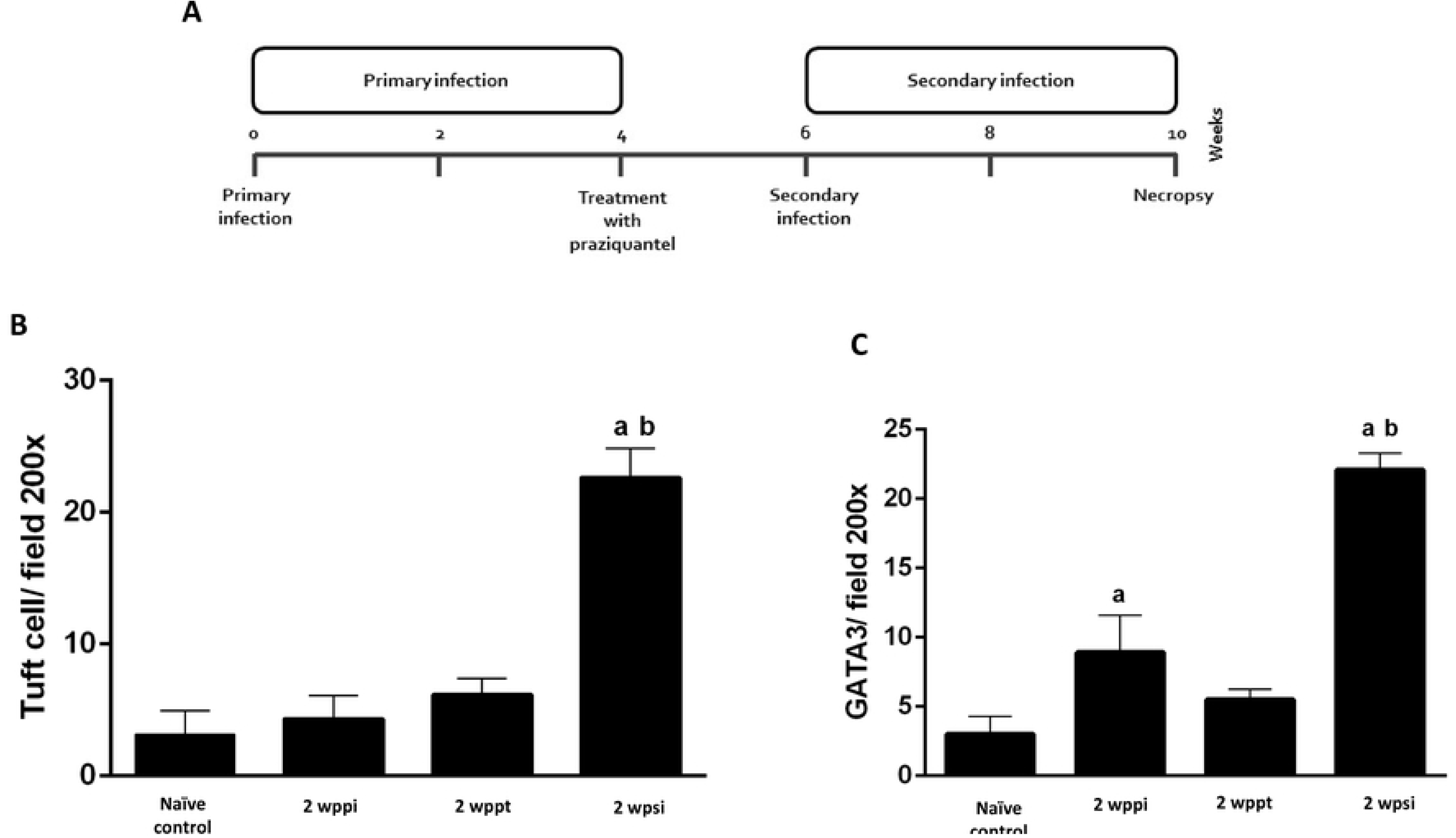
Secondary *E. caproni* infection induces expansion of tuft cells and GATA3+ cells. (A) Schematic representation of the experimental protocol; (B) Counts of tuft cell populations and (C) GATA3+ cells two weeks after primary infection (2wppi), two weeks after treatment with praziquantel (2wppt) and two weeks after secondary infection with *E. caproni* (2 wpsi).

### Blocking of IL-25 reverts resistance against *E. caproni* challenge infections

To analyze the effect of IL-25 in resistance to *E. caproni* challenge infections, a total of 10 mice were given a primary infection and treated with praziquantel at 4 wpi. A total of 5 of those mice were treated with mα-IL-25 before a challenge *E. caproni* infection at 2 wptt. The remainder mice were given a secondary infection at the same time without mα-IL-25 treatment. All mice were necropsied at 2 wpsi.

The results obtained show that blockade of the IL-25 reverted the partial resistance to infection and the number of worms recovered was significantly higher in the animals treated with mα-IL-25 than in non-treated mice (Fig. 4A). Worm recoveries in mα-IL-25-treated mice ranged from 78-89% (84.00 ± 3.9), whereas the values ranged from 9-32% (23.12 ± 8.2) in non-treated animals.

**Figure 4.**
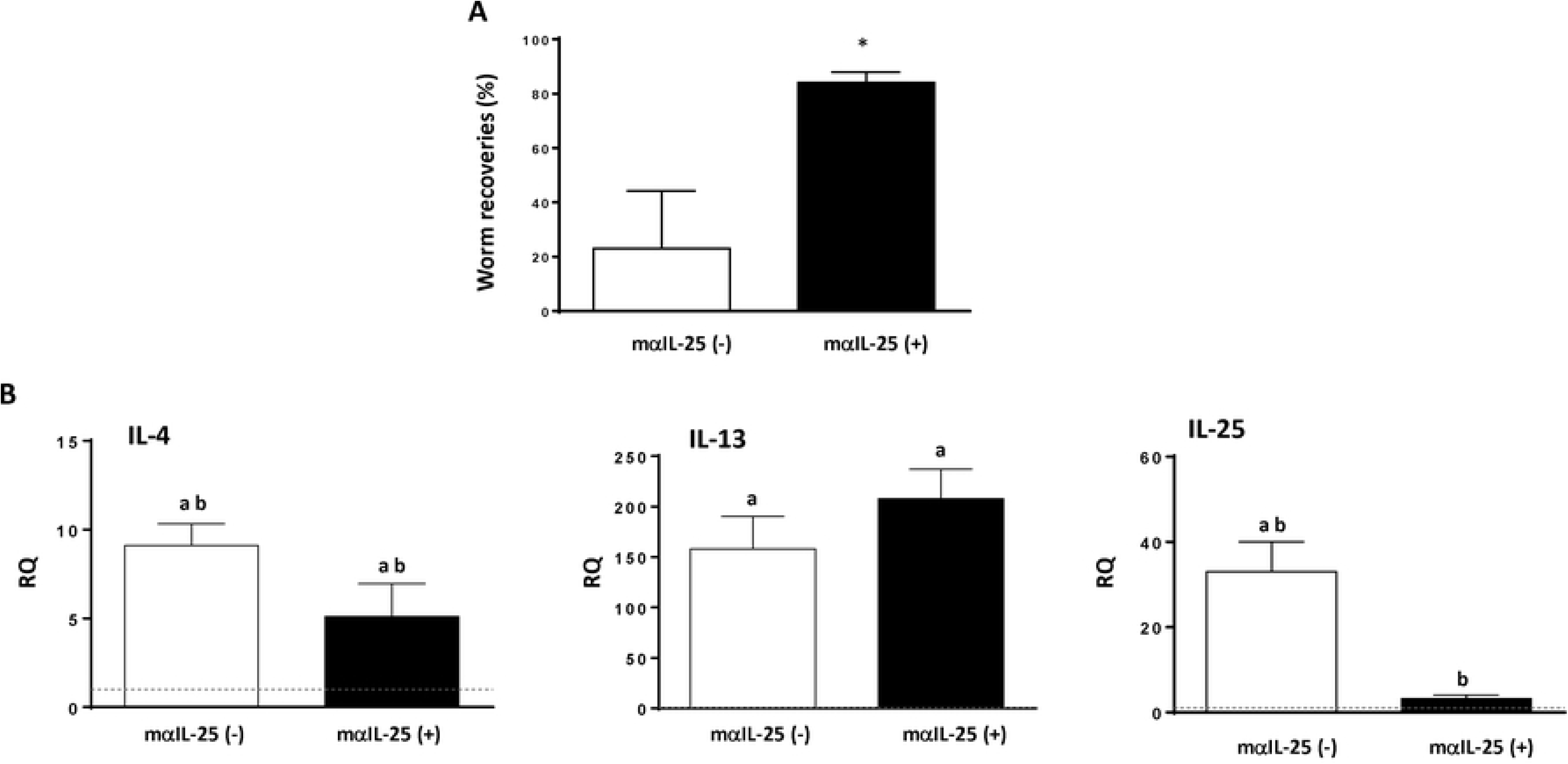
Secondary *E. caproni* infection induces expansion of GATA3+ cells group 2 innate lymphoid cells (ILC2). Kinetics of ILC2 populations two weeks after primary infection (2wppi), two weeks post-treatment with praziquantel (2wppt) and and two weeks after the secondary infection with *E. caproni* (2 wpsi). (A) Immunohistochemical images showing changes in GATA3+ cell populations and (B) ILC2 counts over the course of the experiment. a: significant differences with respect to naïve mice controls; b: significant differences between groups (p<0.05). Scale bar: 30 μm.

Changes in cytokine expression in secondary infection at 2 wpsi in relation to the blockade of IL-25 were investigated by real-time PCR. The most relevant alterations affected IL-4, IL-13 and endogenous IL-25 (Fig. 4B). The remaining cytokines did not show significant differences between treated and non-treated animals or with respect to the negative controls (data not shown).

Animals treated with mα-IL-25 showed a similar cytokine profile to non-treated mice. In both groups a Type 2 response was generated with elevated levels of IL-4 and IL-13 expression after the challenge infection. Probably, the most striking feature observed was the significantly lower levels of endogenous IL-25 expression in mα-IL-25-treated animals with respect to non-treated mice (Fig. 4B).

To study the changes in macrophage activation induced by the blockade of IL-25, we have analyzed several markers of classical or M1 (Arg II and iNOS) and alternative or M2 (Arg 1) activation. Interestingly, antibody blockade of endogenous IL-25 did not change the predominance of M2 activation. However, a significant decrease in the expression of Arg I, Arg II and iNOS was observed with respect to the group of non-treated mice (Fig. 5A). Moreover, blocking of IL-25 did not induce changes in RELM-β expression (Fig. 5B).

**Figure 5.**
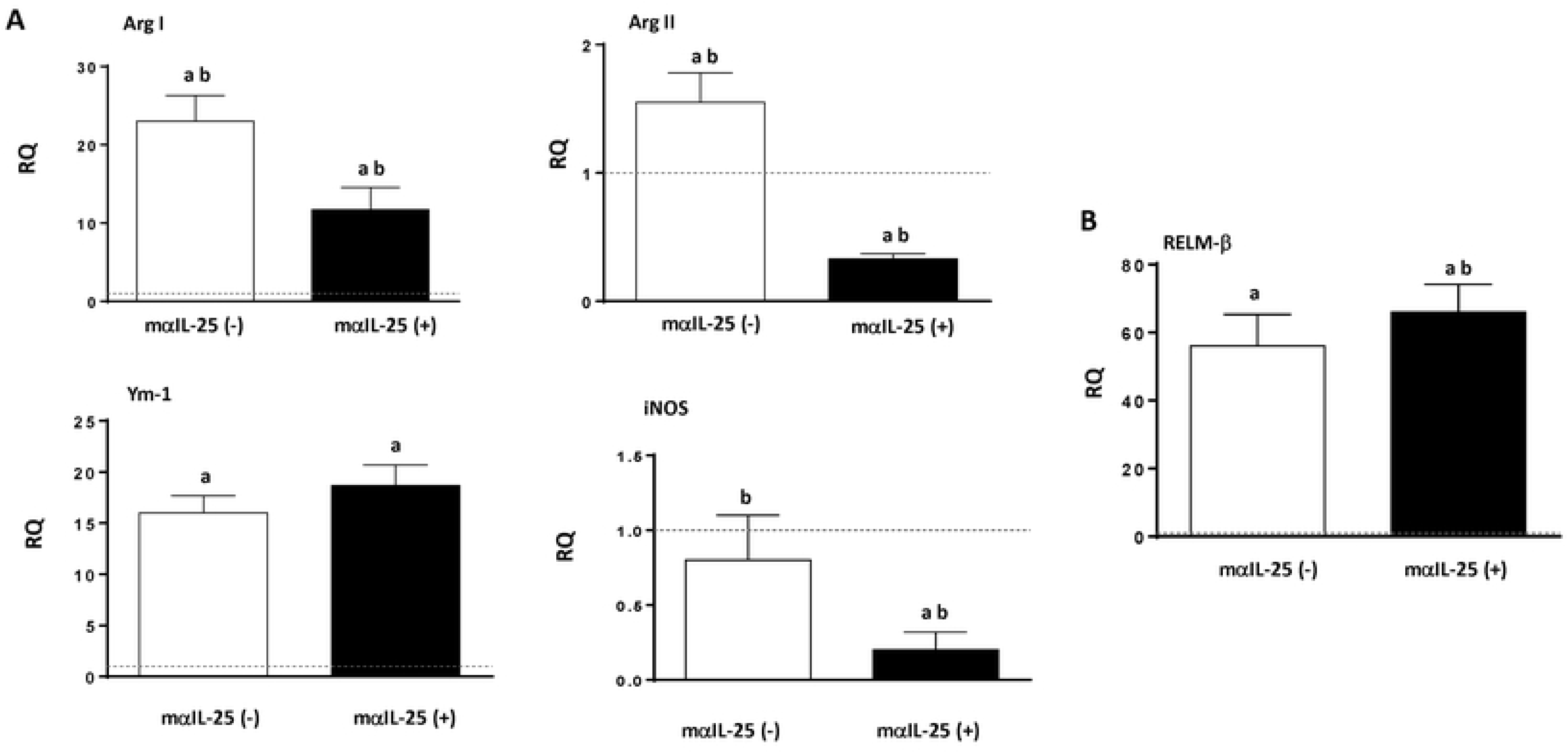
Blocking of IL-25 in challenge infections with *Echinostoma caproni* reverts resistance to infection despite the development of a Th2 response. (A) worm recovery in mα-IL-25-treated ICR mice and non-treated mice after a challenge infection with *E. caproni*; (B) expression of cytokine mRNA in the intestinal tissue of mα-IL-25-treated ICR mice and non-treated mice after a challenge infection with *E. caproni*. The relative quantities (RQ) of cytokine genes are shown after normalization with β-actin and standardization of the relative amount against day 0 sample. Vertical bars represent the standard deviation. *: significant differences with respect to non-mα-IL-25-treated mice. a: significant differences with respect to naïve mice controls; b: significant differences between groups (p<0.05).

### Return to baseline expression of IL-25 after healing of the primary infection ablated the resistance against challenge infection

To analyze further the role of the innately produced IL-25 and the ability of *E. caproni* to induce IL-25 expression in mice in memory secondary infections, we delayed the challenge infection until IL-25 expression recovered to baseline levels. To this purpose, mice were primarily infected with metacercariae of *E. caproni* and treated with praziquantel at 4 wpi. From 6 wppi, goups of those mice were sacrificed every two weeks and the levels of IL-25 expression were studied by rtPCR. Baseline levels were recovered at 10 wppt (Fig. 6). At 10 wppt, the remainder mice of the group were secondarily infected and necropsied at 12 wppt. Moreover, a total of 5 naïve mice of the same age were primarily infected at the same time and used as control of the experiment.

**Figure 6.**
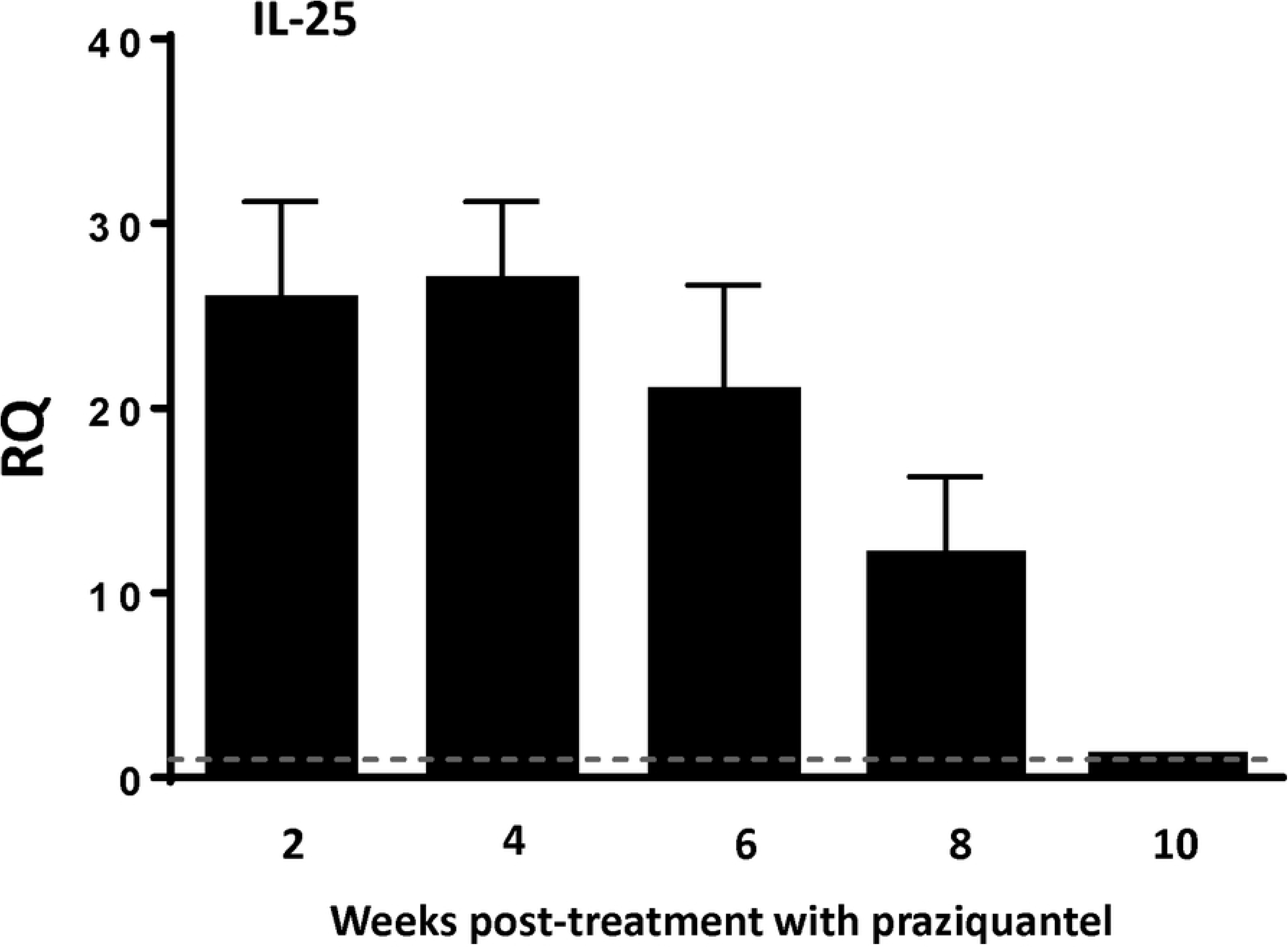
IL-25 expression returned to baseline levels 10 weeks after pharmacological curation of a primary infection. Expression of IL-25 mRNA in the intestinal tissue mice after curation with praziquantel of a primary infection with *E. caproni*. The relative quantities (RQ) of cytokine genes are shown after normalization with β-actin and standardization of the relative amount against day 0 sample. Vertical bars represent the standard deviation. **Blocking of IL-25 induced alternative activation of macrophages after challenge infection.** (A) Pattern of macrophage activation is different in primary and secondary infections analyzed by the expression of markers mRNA of both classical (Arg II and iNOS) and alternative (Arg I and Ym-1) activation of macrophages in in the intestinal tissue of mα-IL-25-treated mice and non-treated mice after a challenge infection with *E. caproni;* (B) expression of RELM-β mRNA in the intestinal tissue of mα-IL-25-treated ICR mice and non-treated mice after a challenge infection with *E. caproni*. The relative quantities (RQ) of cytokine genes are shown after normalization with β-actin and standardization of the relative amount against day 0 sample. Vertical bars represent the standard deviation. a: significant differences with respect to naïve mice controls; b: significant differences between groups (p<0.05).

Results of worm recovery show that after declining levels of IL-25, mice were susceptible to infection again. The worm recovery in secondary infections in absence of IL-25 [46-59% (54.2 ± 11.1)] was similar to that of observed in the animals of the same age primarily infected [49-61% (58.6 ± 13.6)] (Fig. 7A). It should be noted that the number of worms collected in both groups was slightly lower than in other experiments, but this appears to be due to an effect the age of the mice.

**Figure 7.**
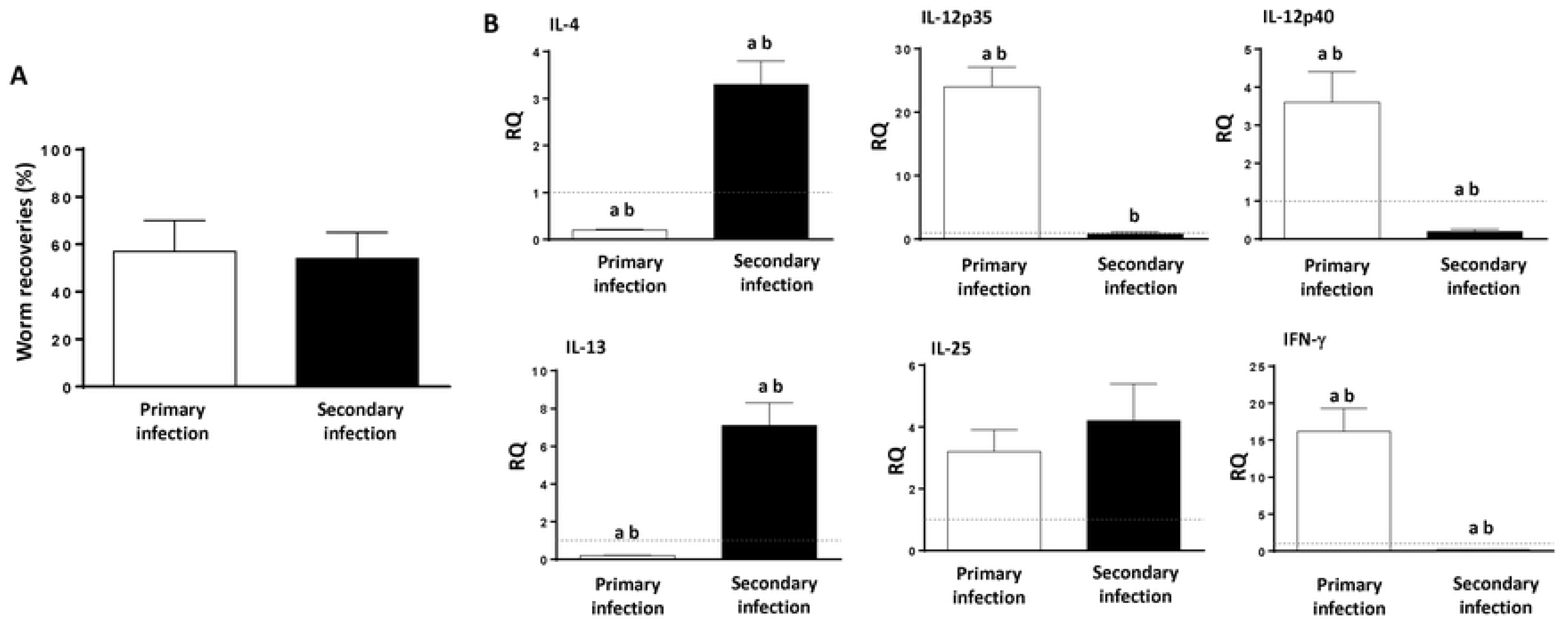
Recovering of baseline expression of IL-25 after healing of the primary infection reverted the resistance against challenge infection together with a Th2 response. (A) worm recovery of a primary infection in naïve mice and in that of a challenge infection in mice in which the basal levels were recovered after the cure of a primary infection; (B) expression of cytokine mRNA in the intestinal tissue of of both groups of mice at 2 weeks after primary and secondary infection, respectively. The relative quantities (RQ) of cytokine genes are shown after normalization with β-actin and standardization of the relative amount against day 0 sample. Vertical bars represent the standard deviation. a: significant differences with respect to naïve mice controls; b: significant differences between groups (p<0.05).215

The profile of cytokine expression showed that secondary infection at 10 wppt induced the development of a Th2 phenotype, despite the lack of IL-25 at the time of infection (Fig. 7B). Secondary infection at 10 wppt was characterized by significant upregulation of IL-4 and IL-13 and unaltered or downregulation of type 1 cytokines such as, IL-12p35 or IL-12p40 or IFN-γ. In contrast, primary infection at this time induced a Th1 response. No differences between both groups was observed in relation to endogenous IL-25 expression (Fig. 7B).

### Activation of STAT6 is not required for resistance against *E. caproni*

To evaluate the role of STAT6 activation in the resistance mediated by IL-25, we used two experimental approaches in resistant rIL-25-treated mice. In a group of mice, we blocked IL-4Rα using monoclonal antibodies before a primary infection with 50 metacercariae. A second group of mice, was treated with rIL-13Rα2 before the infection. The remaining rIL-25-treated mice were exclusively infected with 50 *E. caproni* metacercariae and were used as control of the effect of treatment with IL-4Rα and/or rIL-13Rα2. All mice were necropsied at 2 wppi.

All the 15 mice were refractory to infection in relation to the treatment with rIL-25. In the two groups of animals, blocked IL-4Rα and rIL-13Rα2-treated mice, a significant reduction in STAT6 phosphorylation and translocation with respect to rIL-25-treated control animals was observed at 2 wppi. In the group of treated with antibodies anti-IL-4Rα, the signal was similar to that observed in naïve animals (Fig. 8).

**Figure 8.**
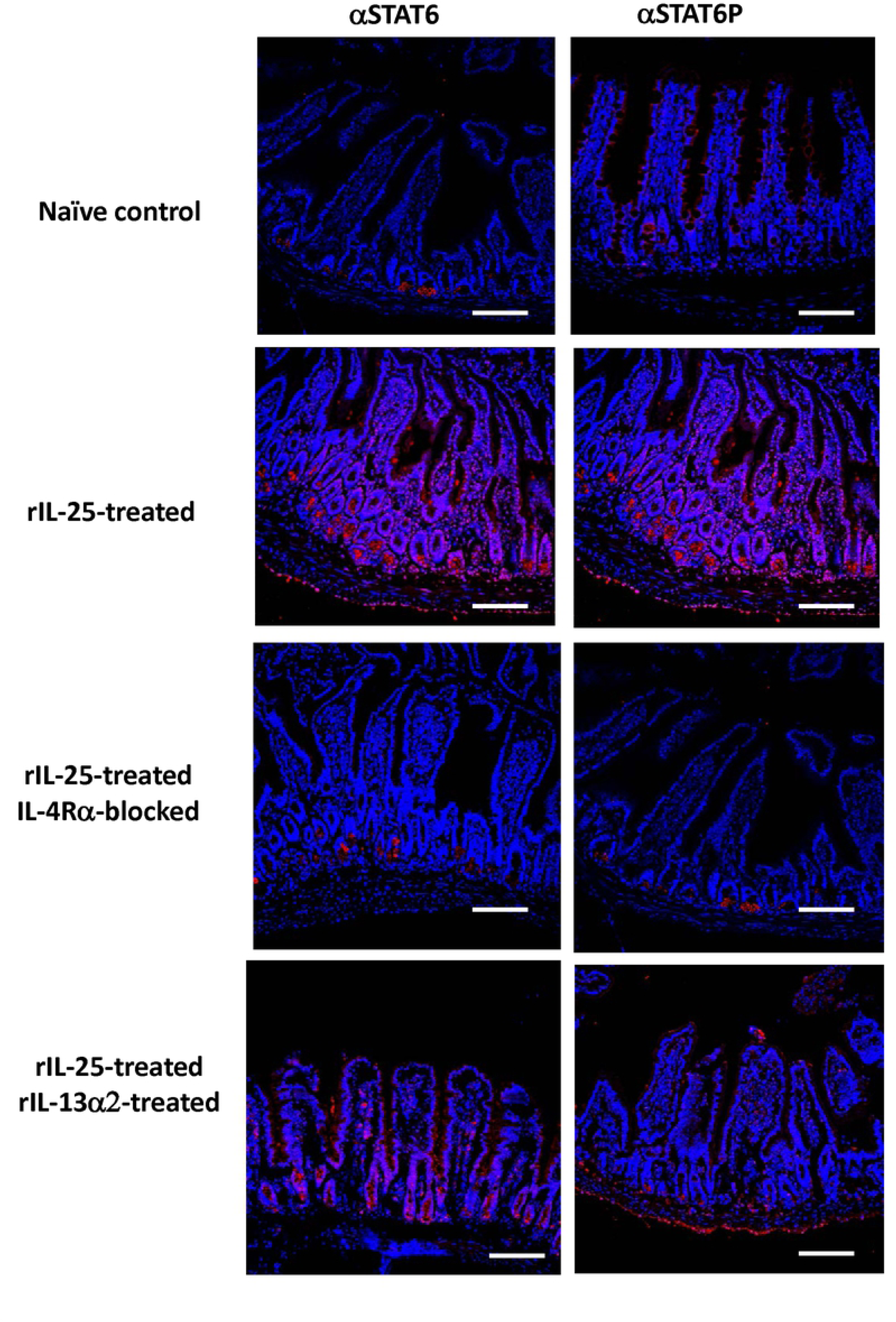
Treatment of IL-25-treated-mice with mα-IL-4Rα or rIL-13Rα2 reduces STAT6 activation. Indirect immunofluorescence with anti-STAT6 (red) and anti-STAT6P (red) on intestinal tissue of IL-25-treated-mice that were also treated with mα-IL-4Rα or rIL-13Rα2 at 2 weeks post-primary infection. Scale bar: 30 μm.

However, the response generated after exposure to metacercariae was different in each group of mice. In the animals lacking IL-4Rα, a type 1 phenotype was developed with elevated levels of IFN-γ expression and no significant changes in IL-4 and IL-13 with respect to naïve controls. No significant changes as compared with naïve mice were observed either in the endogenous expression of IL-25 or in the other cytokines studied. In contrast, treatment of mice with rIL-13Rα2 abrogated any response to infection and no increase in the expression of any cytokine was observed. A decline in the expression of IL-4 and endogenous IL-25 with respect to naïve mice was observed (Fig. 9). The expression of the remaining cytokines remained unaltered in this group of mice. Moreover, slight increases in the expression of iNOS and RELM-β were observed in the group of mice treated with anti-IL-4Rα (Fig 10).

**Figure 9.**
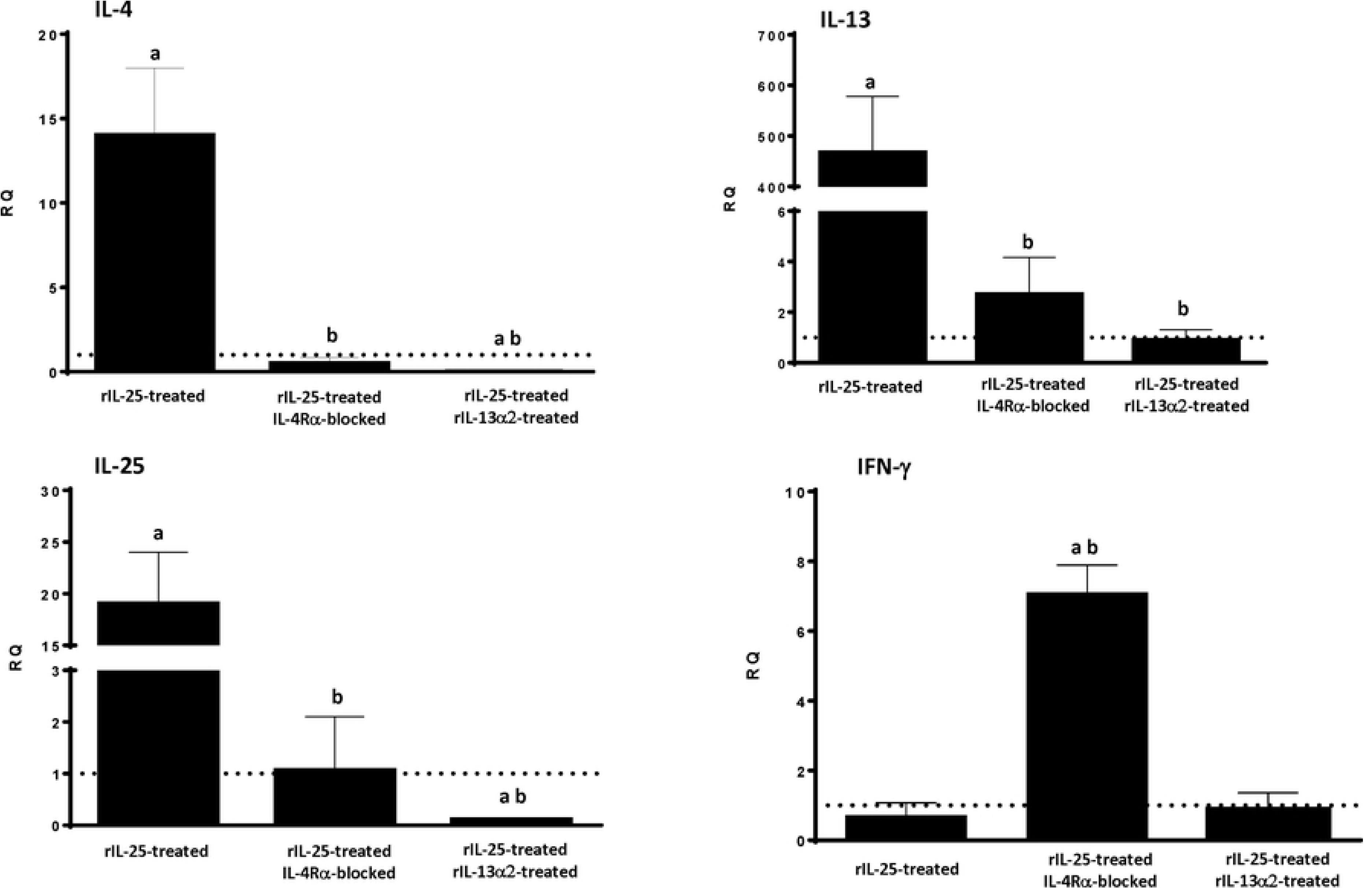
Treatment of mice with either mα-IL-4Rα or rIL-13Rα2 abrogates Th2 response despite the presence of IL-25. Expression of cytokine mRNA in the intestinal tissue IL-25-treated-mice that were also treated with mα-IL-4Rα or rIL-13Rα2 at 2 weeks post-primary infection with *Echinostoma caproni*. The relative quantities (RQ) of cytokine genes are shown after normalization with β-actin and standardization of the relative amount against day 0 sample. Vertical bars represent the standard deviation. a: significant differences with respect to naïve mice controls; b: significant differences between groups (p<0.05).

**Figure 10.**
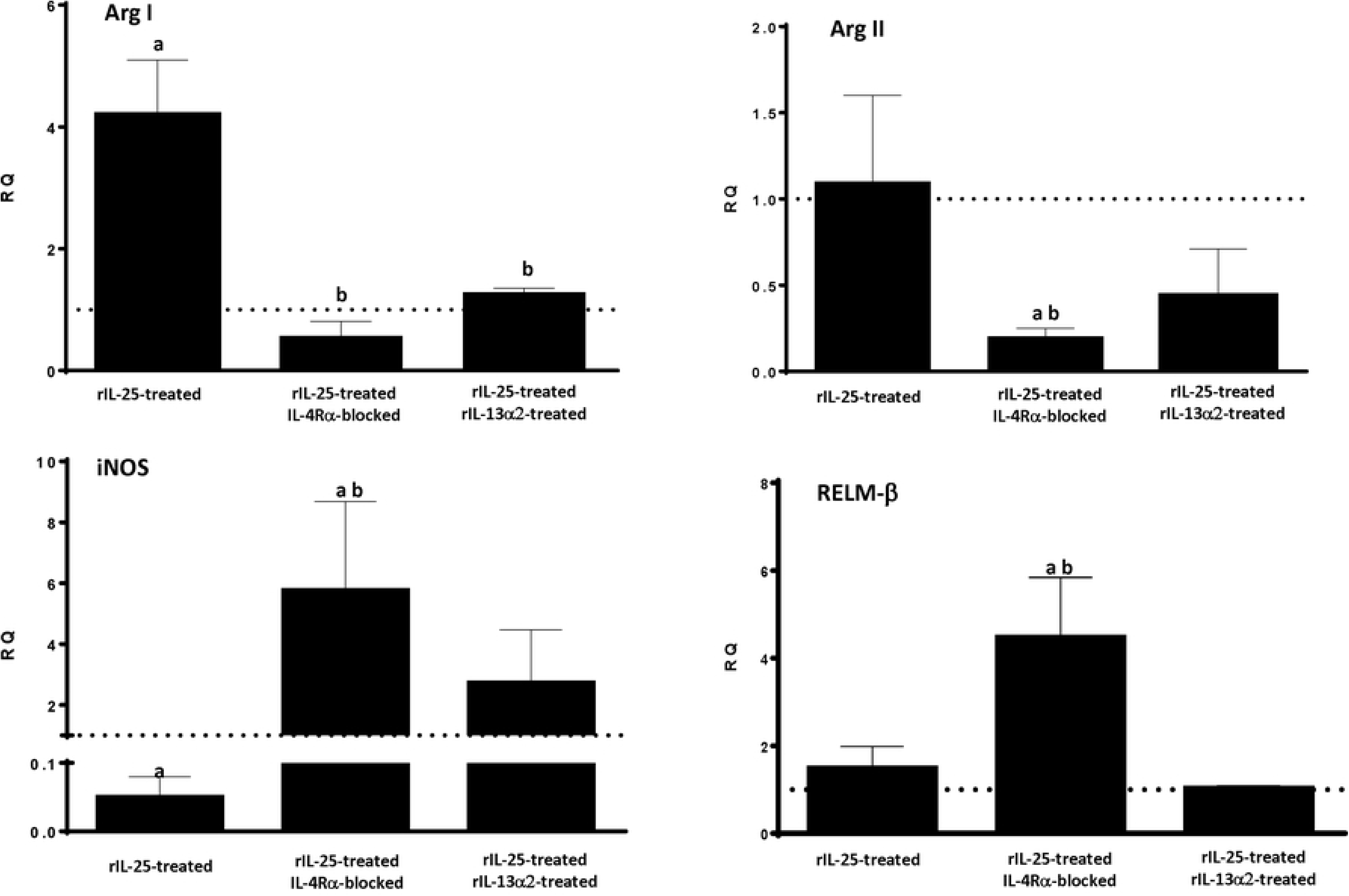
Treatment of mice with either mα-IL-4Rα or rIL-13Rα2 alters the pattern of macrophage activation despite the presence of IL-25. Pattern of macrophage activation analyzed by the expression of markers mRNA of both classical (Arg II and iNOS) and alternative (Arg I and Ym-1) activation of macrophages in the intestinal tissue of IL-25-treated-mice that were also treated with mα-IL-4Rα or rIL-13Rα2 at 2 weeks post-primary infection with *Echinostoma caproni*. The relative quantities (RQ) of cytokine genes are shown after normalization with β-actin and standardization of the relative amount against day 0 sample. Vertical bars represent the standard deviation. a: significant differences with respect to naïve mice controls; b: significant differences between groups (p<0.05).

The expression of IL-13Rα2 in different situations was studied by quantitative PCR. Primary infection resulted in a slight increase in the expression, which increased greatly after pzq-treatment and just before secondary infection associated with resistance. expression declined to negative values after secondary infections (Fig. 11).

**Figure 11.**
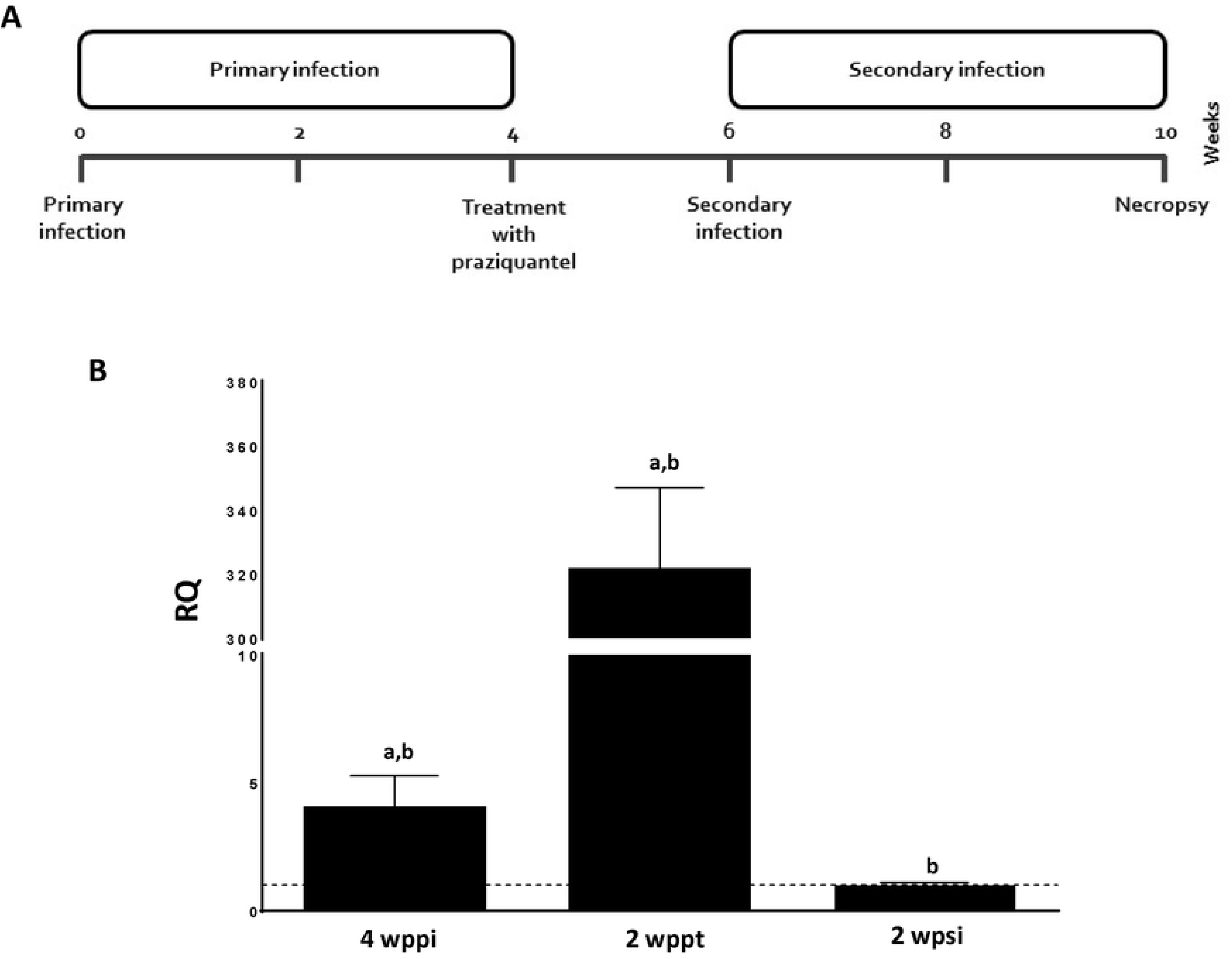
Pharmacological curation of an *Echinostoma caproni* primary infection exacerbates the expression of IL-13Ra2. (A) schematic representation of the experimental protocol; (B) Levels of expression of IL-13Ra2 in the intestinal tissue of mice primarily infected at 4 weeks post infection (4wpi), at 2 weeks post-treatment with praziquantel (2wppt) and at 2 weeks post-secondary infection (2 wpsi) The relative quantities (RQ) of cytokine genes are shown after normalization with β-actin and standardization of the relative amount against day 0 sample. Vertical bars represent the standard deviation. a: significant differences with respect to naïve mice controls; b: significant differences between groups (p<0.05).

### IL-25 production appears to depend on the resident microbiota

To analyze the influence of intestinal resident microbiota in the non-specific upregulation of IL-25 after cure of a primary infection, we initially analyzed by RT-qPCR the changes in the bacterial load as a consequence of the infection. Results obtained showed that *E. caproni* primary infection induced a significant quantitative reduction of bacterial load. After pzq treatment and cure of the primary infection the bacterial charge was recovered (Fig.12A).

To further confirm the involvement of intestinal microbiota in the regulation of IL-25 expression, we treated with a cocktail of broad spectrum antibiotics to maintain the dysbacteriosis a group of mice from 4 wppi (immediately after treatment with praziquantel) to 6 wppi, when they were secondarily infected. The results were compared with a group of mice that were infected, treated with pzq at 4 wppi and challenged at 6 wppi. All mice were sacrificed at 8 wppi. Analysis by PCR showed that antibiotic-treated mice did not produce IL-25 in response to secondary *E. caproni* infection. In contrast, non-treated mice responded with elevated levels of IL-25 expression to secondary infection (Fig. 12B).

**Figure 12.**
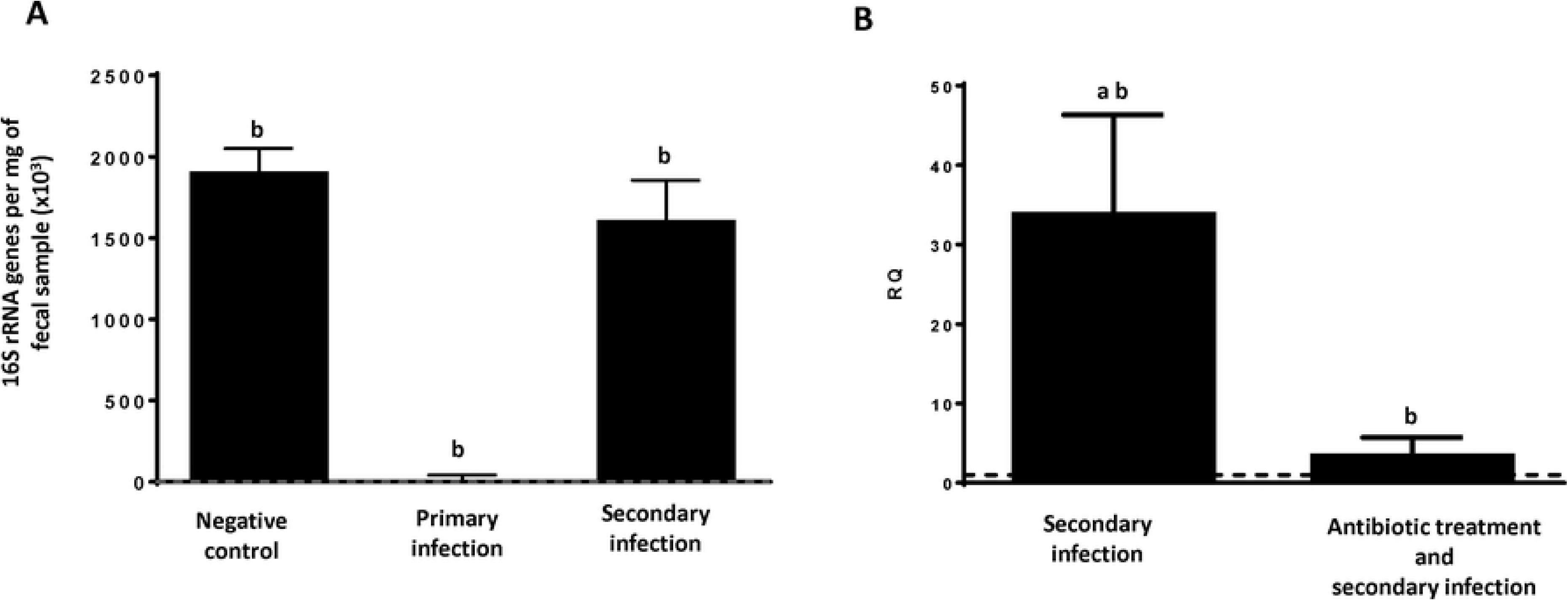
Changes in quantitative composition of resident microbiota alters the production of IL-25 after *Echinostoma caproni* infection. (A) quantitative evolution of resident microbiota as a consequence of a primary and secondary infection with Echinostoma caproni analyzed by quantitative PCR of the 16S rRNA gene of *Escherichia coli* DH5a strain and expressed as 16S rRNA genes per mg of fecal sample (x103). (B) Expression of IL-25 mRNA in the intestinal tissue of secondarily infected mice with or without previous antibiotic treatment. The relative quantities (RQ) of cytokine genes are shown after normalization with β-actin and standardization of the relative amount against day 0 sample. Vertical bars represent the standard deviation. a: significant differences with respect to naïve mice controls; b: significant differences between groups (p<0.05).

## Discussion

Primary infection of mice with *E. caproni* does not elicit IL-25 upregulation. However, pharmacological curation of the primary infection induced a marked overexpression of IL-25. Due to these elevated levels of IL-25, mice became resistant to a challenge infection at 2 wppt, concomitantly with the development of a robust Th2 response [26–27]. Extending our previous studies, the present work was designed to analyze the role of IL-25 in the generation resistance to *E. caproni* infections in mice. The study has been focused on the factors determining IL-25 upregulation, the immune regulatory role of IL-25 and the effector mechanisms induced by IL-25 that determine resistance.

Although it is generally accepted that IL-25 is critical for resistance against intestinal helminths, there is not consensus in relation to the mechanisms by which this cytokine enhances resistance. Traditionally, the role of IL-25 in resistance has been exclusively attributed to its immune regulatory activity. IL-25 promotes Th2 immunity with production of IL-4 and/or IL-13 which, in turn, induce STAT6-mediated intestinal alterations determining parasite rejection [9,31–34]. Upon helminth establishment, intestinal tuft cell populations expand and not specifically release IL-25 that activate a variety of immune cells to initiate type 2 responses and promote Th2-cell-mediated immunity. In response to IL-25 and other alarmins, ILC2 produce large amounts of IL-13 that polarize naïve CD4+ T cells into Th2. Antigen-presenting cells, such as basophils and dendritic cells, are also activated and induce Th2 polarization through different mechanisms [6,13,15]. This is consistent with our results, since we observed that in resistant secondary infections against *E. caproni* there were an expansion of the populations of tuft cells and GATA3+ cells. The effector mechanisms activated in the resistance response against *E. caproni* seem to be dependent on the expansion of the tuft cells, thereby promoting the overexpression of IL-25 and activation of ILC2 or Th2 cells. In an environment of helminth infection, IL-25 acts as a mediator of the activation of ILC2s promoting the polarization of the immune response towards a Th2 phenotype [35].

A number of laboratories have reported that IL-25 upregulation is induced by the infection with intestinal helminths such as *Nippostrongylus brasiliensis, Trichinella spiralis, Trichuris muris* or *Heligmosomoides polygyrus* leading to activation of type 2 responses and resistance to infection [9,31–33,36]. Despite these studies, recent works have suggested that the regulatory role of IL-25 may be secondary and this cytokine operates autonomously from Th2 response in the generation of resistance against intestinal helminths [8, 16]. Smith and co-workers [8] showed that IL-25 plays a more important role than simply the promotion of protective Th2 responses. In fact, these authors demonstrated that adaptive Th2 response to *H. polygyrus* in mice developed normally even in the absence of IL-25R activation, but effector mechanisms became impaired. Similarly, Mearns et al. [16] challenged the role of IL-25 in the promotion of Th2 responses in intestinal helminth infections. Using crossed IL-25^-/-^ C57BL/6 mice and 64 IL-4 C57BL/6 reporter mice, these authors demonstrated no physiological role for IL-25 for either the differentiation of Th2 cells or their development to effector or memory Th2-cell subsets. For instance, IL-25 deficient mice mounted normal Th2 responses following *N. brasiliensis* infections. Our results, indicate that involvement of IL-25 in intestinal helminth infections may be more complex than previously expected. IL-25 is required for resistance against *E. caproni* infection in mice, but this resistance is independent of IL-4 and/or IL-13 activity and STAT6 activation. Furthermore, IL-25 may have a role in promoting Th2 responses though its contribution is different in primary and memory secondary responses. IL-25 is required for the development of a Th2 phenotype in response to primary infections but, in contrast, memory response is characterized by the upregulation of type 2 cytokines despite the lack of IL-25, suggesting that IL-25 enhances the expansion memory cells.

The inability of mice to produce IL-25 in response to primary infection and, consequently IL-13, results in susceptibility to infection. However, treatment of mice with rIL-25 induced resistance to infection, concomitantly with elevated levels of IL-13 [26]. Our results confirm that the development of a Th2 response relies on the presence of IL-13 and STAT6 activation. Despite the lack of IL-25, treatment of mice with rIL-4 or rIL-13 elicited a Th2 phenotype in response to *E. caproni* primary infection and the activation of several IL-13-mediated mechanisms such as goblet cell hyperplasia or RELM-β activation. Our results also support that STAT6 activation is required for the production of type 2 cytokines in response to *E. caproni* primary infection. Blocking of IL-4Rα in mice treated with rIL-25 induced a decline in STAT6 phosphorylation and a Th1 response to infection. Furthermore, our results suggest that IL-13Rα2 plays an important role in the regulation of the response to primary infection. Treatment of mice with IL-13Rα2 abrogated the immune response to *E. caproni* primary infection despite the presence of rIL-25.

IL-4 and IL-13 share a common receptor, the IL-4Rα chain, but IL-13 also uses IL-13Rα1 for signaling via JAK1 and JAK2. IL-13 binds 13Rα1 which complexes with IL-4Rα to form the type 1 receptor signaling, but IL-13 also binds the cell surface and soluble forms of the monomeric type 2 receptor (IL-13Rα2 chain). However, IL-13Rα2 has a decoy effect, lacking signal transduction machinery and limiting the activity of IL-13 since binds the cytokine making it unavailable for activating type 1 receptor [37–40]. Herein, we have shown that rIL-13Rα2 chain limits the ability of mice to respond to *E. caproni* primary infection, even in the presence of rIL-25. Treatment of mice with both rIL-25 and rIL-13Rα2 abrogated the response to *E. caproni* infection and no changes in cytokine levels were observed as a consequence of the infection. IL-13Rα2 may act as negative regulator of both IL-13 inhibiting signal transduction and STAT6 activation by the preferential binding of IL-13 to IL-13Rα2 [41]. However, IL-13Rα2 also inhibits IL-4 induced STAT6 activation and interact with IL-4Rα, even in the absence of IL-13. IL-13Rα2 probably blocks the activation of STAT6 by the physical interaction between the short domain of with the cytoplasmic domain of the IL-4Rα chain that harbors the STAT6 docking sites [37,40–41]. In fact, we have observed that IL-13Rα2 expression if importantly upregulated coinciding with the resistance to secondary infection at 2 wppt.

In contrast to which occurs in primary infections, IL-25 does not appears to be required for the development of Th2 responses in secondary *E. caproni* infections. Herein, we have shown that mice also are unable to produce IL-25 in a secondary challenge infection. Despite this fact, secondary *E. caproni* infection at 2 wppt induced a Th2 response. This was attributed to the presence of elevated levels of IL-25 produced after the cure of the primary infection [26]. However, blocking of the IL-25 innately produced after healing of the primary infection gave rise to a type 2 response as a consequence of the secondary infection showing that IL-25 was not related to the biasing of the immune response. This is in contrast with the results obtained with other intestinal helminths. In *H. polygyrus* infections, both primary and secondary infections included IL-25, but both responses were different. IL-25 response in secondary infections was higher, concomitantly with a more potent Th2 response and enhanced resistance to infection. In contrast, the lower levels of IL-25 overexpression to primary infections, was reflected in a weak response of Th2 cytokines and chronic infections [36]. Although IL-25 does not appear to determine the polarization of Th2 cells in secondary *E. caproni* infections, it may critical for the development to memory Th2 cell subsets. Mearns and co-workers [16] reported that there was not requirement for IL-25 in the development of Th2 cells during *H. polygyrus* infections. To analyze the role of IL-25 in the generation of memory responses against resistance to *E. caproni*, we delayed the challenge infection until the levels of innate IL-25 upregulation declined to baseline, which occurred at 10 wppt. Mice were susceptible to the challenge infection despite the development of a Th2 phenotype with elevated levels of IL-4 and IL-13 but low levels of endogenous expression of IL-25. This suggest that innately produced IL-25 after healing of a primary infection is involved in the differentiation of memory cells.

Resistance to *E. caproni* primary infection was associated with IL-4-independent mechanisms and based on IL-13 activity and STAT6 activation [24–25]. However, recent studies have suggested that mechanisms of resistance to intestinal helminth infections mediated by IL-25 are not dependent on IL-4 and/or IL-13 activity (Smith et al., 2018). Our results support the notion that IL-25 operates autonomously from type 2 cytokines and the generation of resistance is exclusively mediated by IL-25. Treatment of mice with rIL-4 or r IL-13 did not provide of resistance to a primary *E. caproni* infection in relation to the lack of IL-25, despite the development of a Th2 response.

Blocking of the IL-13 receptors induced a significant reduction of STAT6 phosphorylation concomitantly with a reduction of goblet cell hyperplasia and downregulation of RELM-β. However, both control animals and those with the blocked IL-13 receptors were refractory to primary infection due to the presence of exogenous IL-25. These facts indicate that resistance is exclusively mediated by IL-25 independently of the presence of IL-13 and STAT6 activation, probably in relation to M2 activation. Blocking of the IL-13 receptors took the values of iNOS expression to almost zero, together with overexpression of Arg1 indicating an increased M2 activation. It is well known that IL-25 induces alternative activation of macrophages and this an important mechanism for parasite rejection [8, 42]. Independently of the presence of IL-13, M2 has been shown to be crucial for immunity against several intestinal helminths, such as *H. polygyrus* [43]. An interesting feature is the upregulation of IL-13 after blocking of its receptors. Smith et al. [8] obtained similar results in rIL-25-treated mice in *H. polygyrus* infections, reporting that M2 may represent a major source of IL-13 and these authors demonstrated that IL-4, in addition with IL-4Rα signaling, is required for M2 activation and parasite elimination. Expression of IL-25R in M2 could be required for the parasite expulsion in the presence of IL-4Rα signaling [8]. Our results support that M2 activation and the subsequent IL-13 overexpression do not depend on type I receptor signaling. Although the IL-4 expression was not very high, this cytokine may well acts via type I receptor signaling enhancing the resistance to *E. caproni*. Strikingly, treatment of mice with rIL-4 did not yield neither resistance nor M2 activation, probably in relation to the lack of IL-25 production. This suggest that IL-4, but not IL-13, might be necessary for resistance along with IL-25.

An striking feature of *E. caproni* infections in mice is that tuft cell hyperplasia and the subsequent IL-25 overexpression exclusively occurs as a consequence of the healing of the infection. Howitt and co-workers [14] suggested that IL-25 upregulation in intestinal helminth infection is initiated during colonization by the recognition of parasite compounds by tuft cells via taste chemosensory pathways. To this purpose, tuft cells possess multiple taste-chemosensory G protein coupled receptors and many of them require the G protein subunit gustducin and the transient receptor potential cation channel subfamily M member 5 (TRMP5) to transduce the signals [44] Howitt et al. [14] reported that disruption of chemosensory signaling by the loss of TRMP5 abrogated the tuft cell expansion and IL-25 upregulation on mice infected with *N. brasiliensis, T. spiralis,* or *H. polygyrus*. Although other mechanisms of immune suppression cannot be discarded [45], the lack of IL-25 expression in both *E. caproni* primary and challenge infections suggests that parasite components do not activate taste chemosensory pathways in tuft cells of mice which explain the susceptibility to both type of infections. The fact that expansion of GATA3+ cells exclusively occurs after a secondary infection in presence of IL-25, may indicate that only the simultaneous combination of signals provided by the parasite and IL-25 are able to induce the polarization to Th2.

Strikingly, the mucosal regeneration and healing processes initiated after deworming appears to be implicated in the signaling leading to tuft ell hyperplasia and IL-25 upregulation. *E. caproni* induces severe epithelial damage in mice and several mechanisms for wound healing are activated early after primary and secondary infection [1-323,29-30,46]. In the murine gut, wound environment induces rapid changes in resident microbiota such as changes in microbial alpha or beta diversity [47]. Several intestinal nematodes such as *N. brasiliensis, Trichuris trichiura or Ascaris lumbricoides* induce significant alterations in diversity and composition of the intestinal microbiota [48–51]. Moreover, changes in microbial composition associated with parasite infections elicit upregulation of cytokines altering the regulation of the immune response [52–54]. Administration of probiotics promoted successful establishment of *H. polygyrus* in mice, via reduction of Th2 cytokines such as IL-4 a and IL-13 and an increase in regulatory TCD4+ cells [55]. In contrast, resistance to *T. spiralis* was enhanced by promoting Th2 responses after oral administration of *Lactobacillus casei* [56]. In the case of IL-25, several studies support that its intestinal production is regulated by resident microbiota showing that, in general, dysbacteriosis upregulates IL-25 expression [57]. Moreover, IL-25-mediated intestinal immune regulation is impaired in mice in absence of microbiota [58–60]. Expression of ileal IL-25 is reduced in in germ-free mice compared to wild-type mice, but exposure to environmental microbes induced IL-25 overexpression [58–59]. Furthermore, antibiotic treatment of mice significantly decreased the expression of gut IL-25 expression [61]. Our results support that expression of IL-25 could be dependent on microbial-derived signals. Changes in resident microbiota as a consequence of the infection and subsequent healing participate in IL-25 production protecting from secondary infection. Treatment of mice with a cocktail of antibiotics abrogated the IL-25 after the curation of the primary *E. caproni* infection concomitantly with a decrease in bacterial abundance in feces and susceptibility to challenge infection at 2 wppt. Changes in resident microbiota may play a pivotal role in the expression of IL-25 and, consequently, in the resistance to challenge infections.

In summary, we have analyzed the role of IL-25 in the resistance against *E. caproni* in both primary and secondary memory responses in ICR mice. Susceptibility of mice relies in the inability of mice to produce IL-25 in response to infection, which is probably related to alterations in the resident microbiota induced by the infection. In contrast to primary infection, secondary infection elicits a type 2 response, even in the absence of IL-25 expression. Despite the development of a type 2 response, mice are susceptible to secondary infection in relation to the lack of IL-25. Resistance to infection is due to IL-25, which acts autonomously from Th2 response in the parasite clearance.

## Acknowledgements

The authors are sincerely grateful to Prof. Rick M. Maizels and Dr Stephan Löser (Wellcome Centre for Integrative Parasitology, Institute of Infection, Immunity and Inflammation, College of Medical, Veterinary, and Life Sciences, University of Glasgow, Glasgow, UK) for the critical reading of the manuscript and the collaboration provided in some of the experiments included in this work. A part of the experiments were performed in the Wellcome Centre for Integrative Parasitology, Institute of Infection, Immunity and Inflammation of the College of Medical, Veterinary, and Life Sciences of the University of Glasgow (Glasgow, UK) by MAI in a stage financially supported by the Erasmus program of the UE and the Cámara de Comercio, Industria, Servicios y Navegacion (Valencia, Spain).

## Legends for figures

**Supplementary Figure 1.** Expression of cytokine mRNA in the intestinal tissue of rIL-4-treated or rIL-13-treated mice. The relative quantities (RQ) of cytokine genes are shown after normalization with β-actin and standardization of the relative amount against day 0 sample. Vertical bars represent the standard deviation. a: significant differences with respect to naïve mice controls; b: significant differences between groups (p<0.05).

**Supplementary Figure 2.** Immunohistochemical images showing changes in tuft cell populations (A) and GATA3+ cells two weeks after primary infection (2wppi), two weeks after treatment with praziquantel (2wppt) and two weeks after secondary infection with *E. caproni* (2 wpsi). Scale bar: 10 μm.

**Supplemetary Table 1** Applied Biosystems inventoried assays used

